# Durability of environment-recruitment relationships in aquatic ecosystems: insights from long-term monitoring in a highly modified estuary and implications for management

**DOI:** 10.1101/256404

**Authors:** Natascia Tamburello, Brendan M. Connors, David Fullerton, Corey C. Phillis

## Abstract

The environment can strongly influence the survival of aquatic organisms and their resulting dynamics. Our understanding of these relationships, typically based on correlations, underpins many contemporary resource management decisions and conservation actions. However, such relationships can break down over time as ecosystems evolve. Even when durable, they may not be very useful for management if they exhibit high variability, context dependency, or non-stationarity. Here, we systematically review the literature to identify trends across environment-recruitment relationships for aquatic taxa from California’s San Francisco Bay and Sacramento-San Joaquin Delta Estuary. This is one of the most heavily modified aquatic ecosystems in North America, and home to numerous species of concern whose relationships with the environment inform regulatory actions and constraints. We retested 23 of these relationships spanning 9 species using data that have accumulated in the years since they were first published (9-40 additional years) to determine whether they persisted. Most relationships were robust (i.e., same or stronger in magnitude) to the addition of new data, but the ability to predict how a species will respond to environmental change did not generally improve with more data. Instead, prediction error generally increased over time and in some cases very quickly, suggesting a rapid regime shift. Our results suggest that more data alone will not necessarily improve the ability of these relationships to inform decision making. We conclude by synthesizing emerging insights from the literature on best practices for the analysis, use, and refinement of environment-recruitment relationships to inform decision making in dynamic ecosystems.

## Introduction

The environment can have a profound and complex influence on aquatic organisms and their population dynamics (e.g., Szuwalski et al. 2015). Understanding when and how the environment influences the survival, abundance, and recruitment of fishes and other aquatic organisms has long fascinated and perplexed fish and fisheries scientists and managers (Hjort 1914; Cushing 1995; Jacobson and MacCall 1995). Knowing how the environment influences the survival and dynamics of fishes is of general ecological interest because of the light it can shed on, for example, the relative influence of bottom-up and top-down control in ecosystems. Quantifying how the environment influences recruitment can in theory help inform fisheries and improve management. Given the now pervasive influence of humans over the world’s aquatic ecosystems (e.g., Halpern et al. 2015), understanding when and how the environment – and human influences on it – affect the dynamics and abundances of aquatic organisms is critical to many decisions in natural resource management.

Relationships between the environment and recruitment, defined here as any relationship between the number of individuals in a population (or their survival rate) and their environment (e.g., river flow, pesticide concentration, water temperature), may break down over time. In a now classic review of fisheries literature, Ransom Myers (1998) found that only 22 of 77 environment-recruitment relationships still held after being re-examined with new data. The relationships that were most likely to stand the test of time were those with temperature at the limit of a species’ range, where the influence of physical tolerance thresholds outweighs that of more complex ecological interactions. Even when such correlations are reliable, they may not be very useful for informing management if the described relationship is characterized by high variability, context dependency, or non-stationarity as is often the case with recruitment data. Nonetheless, relationships between the environment (which we define broadly as both natural and those aspects under human control) and fish recruitment are central to contemporary resource management decision-making and the conservation of aquatic species. Considering this, and the potential for environment-recruitment relationships to break down over time, there is a pressing need for guidance on best practices for the analysis, use, and refinement of environment-recruitment relationships to inform decision making in natural resource management.

California’s San Francisco Bay and Sacramento-San Joaquin Delta Estuary (hereafter “Bay Delta”) (Figure 1) is an ideal system in which to examine the durability and usefulness (i.e., predictive power) of environment-recruitment relationships and their implications for decision making. The Bay Delta system has been continuously monitored in a systematic manner for long periods of time, it is heavily altered and managed, and it is home to numerous endangered species and associated regulatory actions and constraints including some which are based upon environment-recruitment relationships (reviewed in Kimmerer 2004). Numerous relationships have been described for taxa in the Bay Delta, but very few have been revisited to test whether these correlations are reliable in the face of new data.

**Figure 1:**
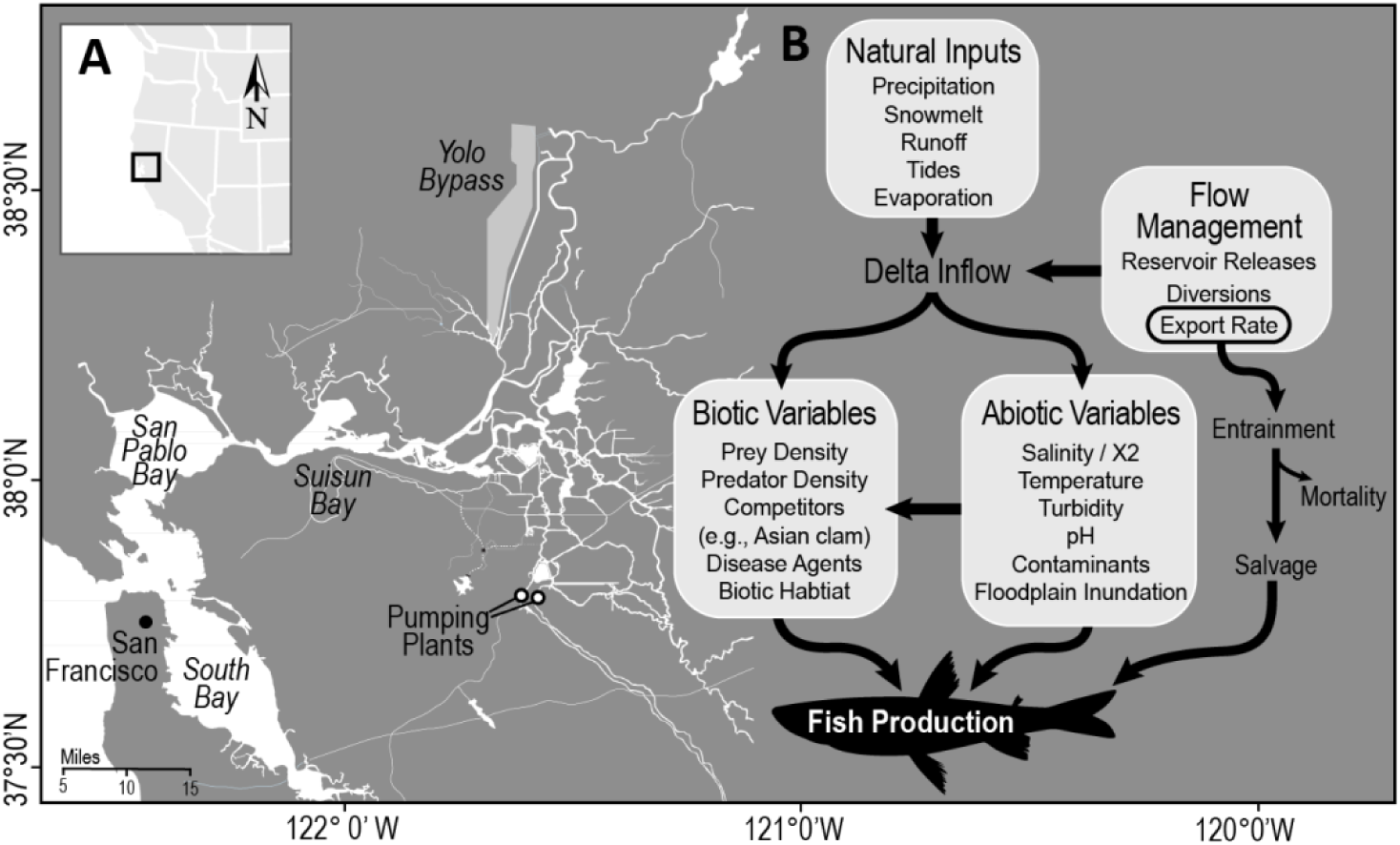
Map of the Bay Delta region and key geographical features (A), along with an overlay of a conceptual model of key abiotic and biotic drivers known to influence fish production in this system (B). Adapted from the Delta Independent Science Board 2015.

Here, we systematically review the literature to identify environment-recruitment relationships for Bay Delta taxa. We then reanalyze the relationships where new data are available to quantify the extent to which the relationships still hold when confronted with new data. Finally, we synthesize emerging insights from the literature on best practices for the analysis, use, and refinement of environment-recruitment relationships to inform decision making in natural resource management.

## Methods

### Study area

The Bay Delta is made up of a large interior delta formed by the Sacramento and San Joaquin Rivers feeding into a series of basins separated by narrow, deep tidal channels, which flow into a seaward region and ultimately into the San Francisco and San Pablo Bays which are connected to the Pacific Ocean (Figure 1A). The Bay Delta is one of the most heavily modified estuaries in the United States, and is strongly influenced by state and federal water project operations. Two pumping facilities export delta inflows to meet metropolitan and agricultural water needs. Water exports can affect fish directly through entrainment into the pumping facilities, and indirectly though the influence of reduced flows on a wide range of abiotic and biotic variables (Figure 1B). Established quantitative relationships between flow characteristics and fish abundance, survival, and migration underpin regulatory decisions regarding levels of allowable exports of river flows and minimum outflow from the Bay Delta that maintain fish production and habitat within acceptable bounds (CDFW 2016a,b).

### Literature review

We carried out a literature search of peer-reviewed publications, grey literature, and government agency reports to identify published examples of relationships between environmental variables and population parameters of aquatic Bay Delta species. Our initial search was carried out using Google Scholar and various combinations of search terms based on variables expected to influence organism abundance as described in comprehensive reviews of Bay Delta ecology (e.g., Kimmerer 2004; CDWR and USBR 2016). These search terms included “fish”, “invertebrate”, “abundance”, “survival”, “entrainment”, “migration”, “environment-recruitment”, “environmental variable”, “X2”, “flow”, “conductivity”, “turbidity”, and “prey density”. The resulting set of publications and reports was then supplemented by consulting regulatory documents to identify additional relationships that underpin contemporary management decisions.

We created a catalogue of all identified publications to document their various characteristics, including publication year, focal species, the number of relationships examined, and whether the publication is cited in regulatory documents as informing management decisions (included here as Supplementary Online Material). Because most of the government agency and grey literature reports were review documents that reproduced results from primary literature, we chose to focus further investigations only on the peer-reviewed literature. For each peer-reviewed study in our catalogue, we extracted each individually reported relationship into a second catalogue and documented characteristics including the focal species, predictor and response variables, type of analysis, timeframe, primary data source, and reported model outputs including intercept and slope parameters, R^2^, p-value, AIC, and others. This catalogue of relationships was used both to carry out a qualitative analysis of trends in the study of such relationships in the Bay Delta, and to select a subset of environment-recruitment relationships suitable to retesting to quantify the extent to which the relationships still hold when confronted with new data.

### Relationship selection criteria

We developed three tiers of criteria used to screen the full set of published environment-recruitment relationships to identify those suitable for reanalysis.

Tier 1 criteria excluded relationships for which reanalysis would be impractical due to (1) inability to obtain new data for reanalysis because of reliance on either one-time experiments (e.g., paired releases of tagged fish), data collection programs that have since ended, or second-order variables derived via complex integration of many other environmental variables; (2) use of analytical methods that would be impractical to replicate for a review study of this scale (e.g., whole ecosystem simulation models) or which make it difficult to compare strength and statistical support across relationships (e.g., non-linear correlations such as GAMs, rank analyses, etc.); or (3) because the published relationship is so recent there would be few additional data points available (i.e., we excluded relationships if the number of years of new data were less than the number in the original time series or less than 10 years, whichever was less). In practice, this resulted in retention of linear correlations based on data from long-term monitoring programs with publicly available data sets, which formed a large majority (~75%) of all the published relationships in our dataset examined prior to the application of screening criteria.

Tier 2 criteria screened out relationships with little or no statistical support in the initial analysis, given that the objective of our work was to evaluate the extent to which established relationships hold over time. For this purpose, we defined little or no statistical support as p > 0.1, or ΔAIC > 2 from the top model or, where neither is reported in older publications, the absence of a fitted regression line and equation on a scatter plot of the relationship.

Lastly, Tier 3 criteria were used to reduce redundancy in the remaining set of relationships by selecting only a single relationship for each unique species – variable pair for reanalysis. When more than one relationship existed for the same species-variable pair relying on the same source data (e.g., delta smelt ~ X2), the following sub-criteria were used to select a single relationship that was: (1) most recently updated among the set to avoid reproducing past retests; (2), the longest time-series among the set for greater statistical power; (3) based on indices of abundance rather than extrapolated abundance estimates to reduce propagation of error, and (4) where all relationships in the set are statistically significant, the relationship with the strongest support based on variation explained (e.g., via R^2^) or other rationale provided by the authors. We considered similar relationships using different source data (e.g., delta smelt_(Fall Midwater Trawl)_ ~ X2 and delta smelt_(Bay Study Midwater Trawl)_ ~ X2) distinct and therefore were retained due to known differences in methodology, target life stage, and conclusions that can be drawn from alternative source surveys.

The full catalogue of original and screened publications is available in the Supplementary Online Materials.

### Data sources

For each relationship retained for reanalysis, we sought to obtain the same data used in the original analysis from the source identified by the authors. In some cases, this data was available directly, and in others, annual means of data needed for analysis were derived from raw data sets using the same or similar methods originally described by the authors, including applying data transformations. In some cases, we were not able to reproduce the data using the methods described by the authors and either used a modified method or excluded the relationship from our analysis (Table 1). Population data for these relationships were derived from the California Department of Fish and Wildlife’s fall midwater trawl (FMWT), summer townet survey (TNS), beach seine surveys, and salvage surveys; the Bay Study otter trawl (Bay OT) and midwater trawl (Bay MWT) surveys; and the U.S. Fish and Wildlife Service’s Chinook salmon trawls. Environmental data was derived primarily from Hutton et al., (2015) and calculations therein for X2, from the California Department of Water Resources DayFlow data portal for flows, from environmental data collected alongside population data as part of the FMWT survey, and in some cases directly from the study authors (Table 1, Supplementary Online Materials).

**Table 1:**
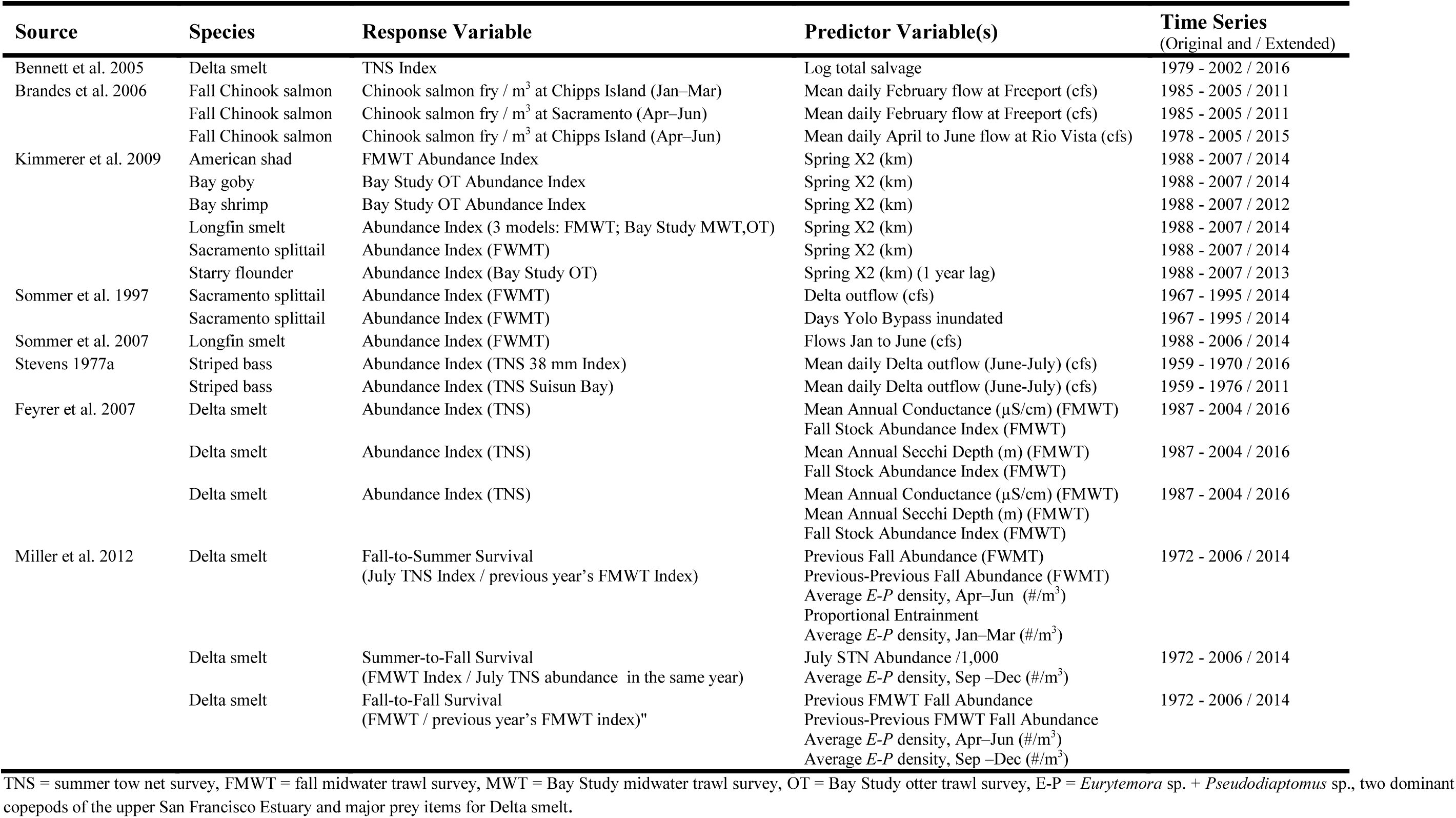
Summary of previously published environment-recruitment relationships retested with additional years of data.

### Analysis

We conducted a quantitative reanalysis of selected environment-recruitment relationships. Source data for the analyses was transformed (as per original analyses; see Table 1), subset to the relevant timeframes, and standardized (by subtracting the mean and dividing by the standard deviation) to allow estimated relationships to be comparable across species and environmental variables. All relationships selected for reanalysis were based on linear correlations and so scaled data was analyzed using ordinary least squares regression with an abundance measure as the response variable and one or more environmental factors as the predictor variable, with none of the ultimately selected relationships including interactions among variables

Several relationships selected for reanalysis (e.g., Kimmerer et al. 2009) included a step-change or data-splitting before and after the introduction of invasive Asian clam (*Corbula amurensis*) into the Bay Delta in 1986. Introduction of this clam is thought to be responsible for a sudden and substantial decline of zooplankton and, subsequently, of fish in the Bay Delta (Kimmerer et al. 1994). Where published analyses found statistical evidence for a split or step change in 1987 (using a dummy variable), we chose to reanalyze the relationship using only data after this cutoff both to simplify the analyses and because prior data could be considered irrelevant for interpreting how relationships have changed in the context of additional data.

For each relationship, we quantified the strength (R^2^; correlation coefficient) and magnitude (slope; in standardized units) of the relationship based on the scaled original and updated time series and then compared them to quantify the extent to which the relationships still held when confronted with new data. We also quantified prediction error using unscaled original and extended time series for each relationship to characterize how well the observed relationships would be expected to predict future (out of sample) observations and to provide an indication of how useful the relationship may be from a decision making and management perspective. Prediction error was estimated as the normalized root mean squared prediction error (*CV*_*n*_):

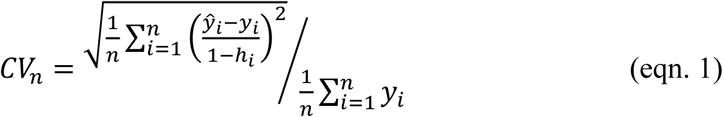

where *h*_*i*_ is the diagonal element of the operator matrix that produces the least squares fit (i.e., hat matrix). This measure of prediction error can be interpreted as the percent error in future predictions relative to the average observed abundance for a given relationship. For example, a prediction error of 100% would mean that the relationship allows us to predict future abundances to within +/- 50% of the mean predicted abundance.

We compared the raw data, parameters, and fit of relationships using the original time series to those reported in the original publications to ensure that our approach successfully replicated the previously published relationships before proceeding to retesting. Instances where we were not able to reproduce past relationships were not further considered (detailed in Table 1).

Our retests of environment-recruitment relationships using data that has accumulated since a relationship was first established has the potential to be biased by an imbalance in the number of pre-to post-retest observations. In instances where there are many (e.g., 30) years of pre-retest observations and only a few (e.g., 10) post-retest observations it is possible that the pre-retest observations obscure what is otherwise a weakening or different relationship in the post-retest observations. To quantify the extent to which this was the case with the relationships we retested, we carried out a secondary analysis where we randomly sub-sampled the pre-retest data so that there was an equal number of pre-and post-retest data points, and then, as above with the full time series, quantified the strength and magnitude of the relationship based on the subsetted data. We repeated this exercise 1000 times for each relationship, and then compared the median strength and magnitude of the relationship based on the original (subsetted) and updated time series.

All analyses were carried out in the R statistical software suite (R Core Team 2017), and we provide the source code, and data, for our analyses in the Supplementary Online Materials.

## Results

### Literature review

Our literature search identified 98 publications describing environment-recruitment relationships in the Bay Delta ecosystem. Of these, 40 were reviews citing primary literature or offered only general observations on raw data without conducting any analysis, and 3 were not available online and so not examined further. The remaining 55 peer-reviewed publications were retained for further analysis. This subset included papers published between 1977 and 2017, with a mean publication date of 2002. These studies examined an average of 10 relationships per study, and a minority of papers examined a large number (>100) of competing models describing the same relationship. Each study examined between 1 and 17 focal species (mean ~ 3).

These 55 peer-reviewed studies described 420 individual relationships which overwhelmingly focused on examining the influence of environmental variables on population abundance as opposed to other biological characteristics (Figure 2A). This is likely a result of the fact that roughly 70% of all relationships relied on publicly available long-term abundance survey data that has been collected in the Bay Delta for decades (e.g., California Department of Fish and Wildlife fall midwater trawl (FMWT), summer townet survey (TNS), and San Francisco Bay Study otter trawl (OT) and midwater trawl (MWT) surveys), whereas other types of population variables are generally not routinely collected for most species and were obtained using one-off experiments or surveys to inform 18% of these relationships. The distribution of relationships was also strongly biased towards species that are either currently or historically listed as Threatened or Endangered (Figure 2B). There was a more even distribution of environmental variables examined across studies, but the most frequently examined variables were X2 (the distance in kilometers from the Golden Gate up the axis of the estuary to where the tidally-averaged near-bottom salinity is 2 ppt or 2 x 10^−9^ mg/L; Jassby et al. 1995), or other measures of salinity, and flow (Figure 2C). Most relationships examined only a single environmental variable (maximum of 7) through simple linear correlation (60% of all relationships). However, the number of variables has generally increased in more recent studies as researchers continue to adopt more complex analytical methods including multivariate models, generalized additive models, and whole-ecosystem simulations. While about 42% of all relationships have been identified as underpinning regulatory decision-making, only 14% of all relationships have been retested to quantify their durability (i.e., extent to which their magnitude of effect holds up in the face of new data).

**Figure 2:**
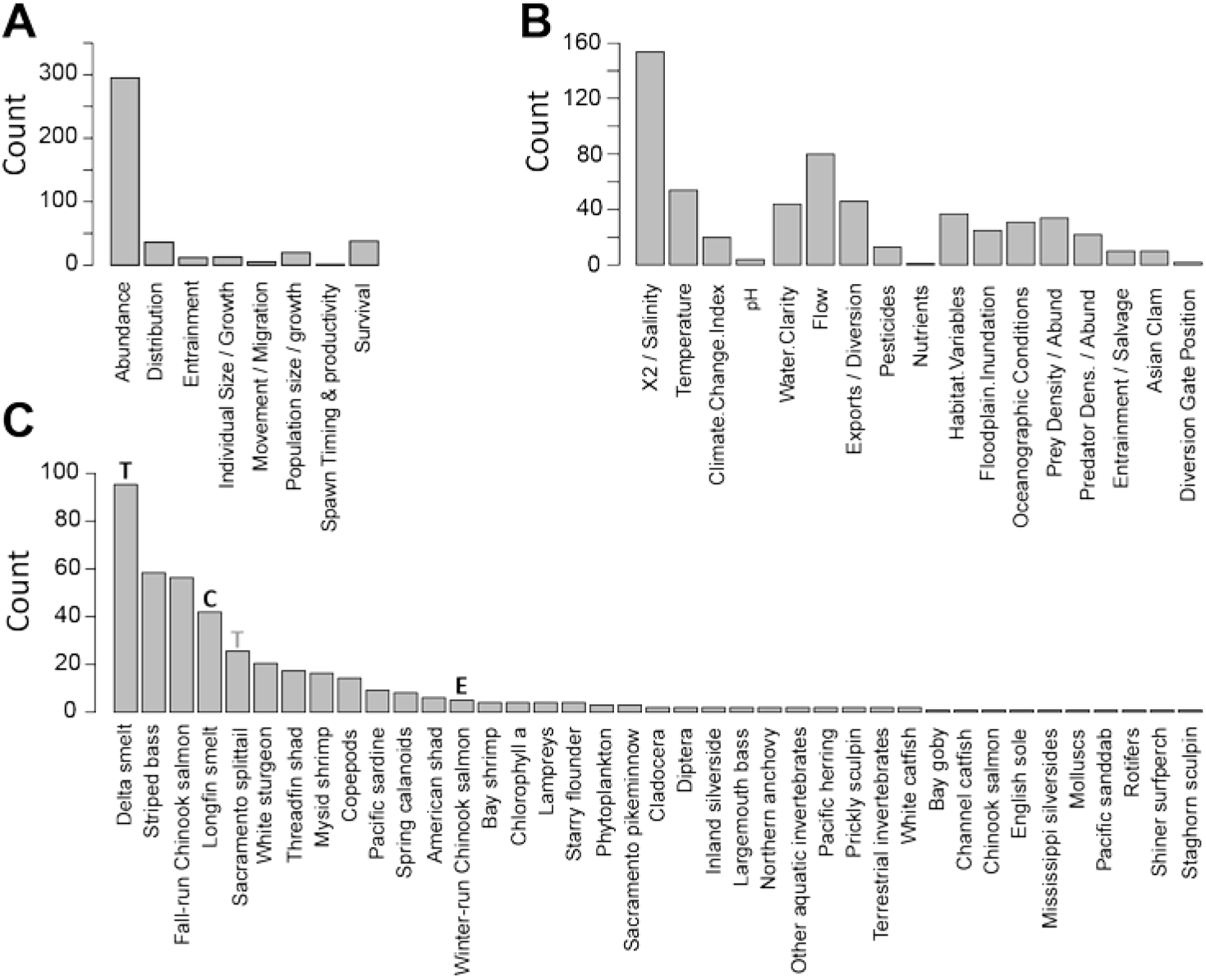
Frequency distributions of the type of response variable (A), environmental variables (B) and focal species (C), represented in the 420 individual environment-recruitment relationships for the Bay Delta region examined in this study, showing the relative emphasis of this body of work. Superscripts in panel C indicate the species current (black) or historical (grey) conservation status as Endangered (E), Threatened (T), or Candidate (C) as reported by the U.S., Fish and Wildlife Service Environmental Conservation Online System.

Across the published relationships there was broad variation within and among environmental variables in the amount of variation in the abundance that was explained. For example, some environmental variables like flow and salinity tended to explain over half the variation, while others such as temperature and water export rates (measured in cfs - cubic feet per second - or m^3^/s, where 35.31 cfs = 1 m^3^/sec) explained much less variation in abundance (Figure 3).

**Figure 3:**
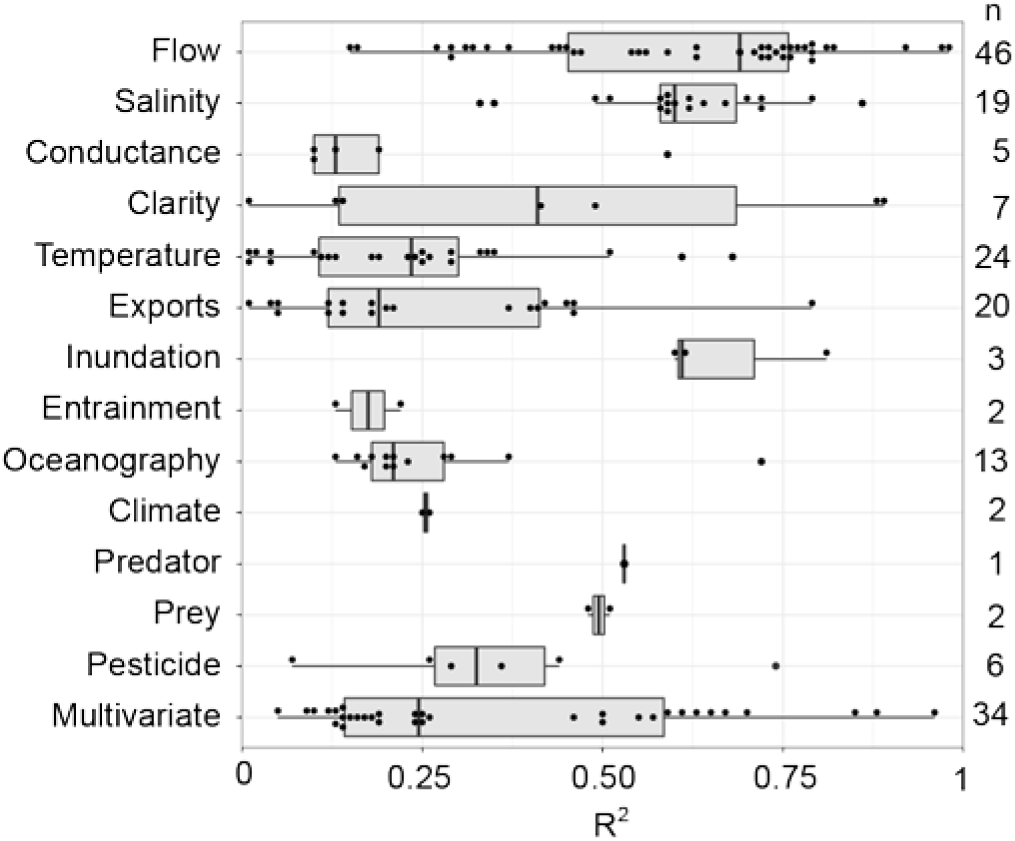
Boxplots and underlying estimates of the distribution of the strength (R^2^) of published relationships for each broad type of environmental variable identified in the literature review (184 papers reported this statistic). Some variables have stronger relationships (e.g., flow, salinity) with the abundance of species than others that are more variable (e.g., temperature, volume of exports) and may therefore depend more on the species and context involved. The “multivariate” environmental variable encompasses relationships that include two or more of the other environmental variables listed here. Black dots outside the range of the box and whiskers represent true outliers (i.e., beyond 1.5 times the interquartile range), while the jittered black dots represent the distribution of remaining data points.

### Reanalysis of bay delta environmental-recruitment relationships

Applying our screening criteria to the full set of relationships resulted in 31 relationships suitable for retesting (Table 1). The publications originally reporting these relationships varied widely in the details provided on methodology, and it was often difficult to determine how the environmental variables were derived or what the source of the data was that was used to derive them. We were unable to complete re-analyses for 8 of these 31 relationships due to an inability to re-create the input variables using the methodology described by the authors or because of errors or missing information in the original publications, leaving 23 relationships (17 univariate and 6 multivariate) from 8 publications which were fully re-analyzed using the most recent available data (between 9 and 40 additional years, median 9 years).

When updated data were used to retest previously published relationships, the direction and statistical significance of the relationships remained the same (Figure 4; Table 1). Of the univariate relationships, 9 of 17 became stronger (i.e., either more negative or positive depending on the original relationship), 3 of 17 became weaker and 5 of 17 remained nearly identical. These general patterns remained the same for the multivariate relationships (Table 1) and when the dataset used to retest relationships was balanced to achieve an equal number of original and updated data points (Figure 4; Table 1).

**Figure 4:**
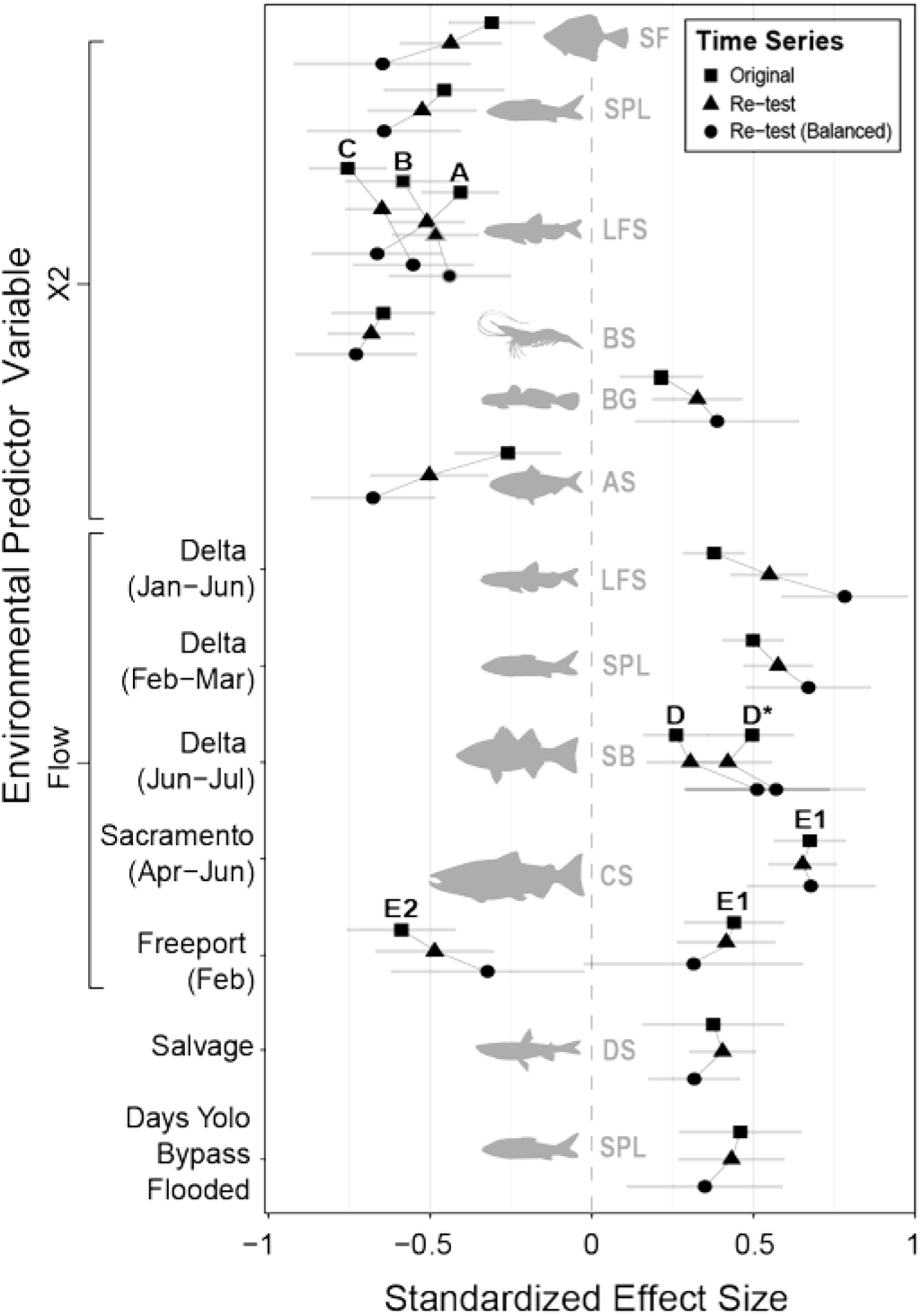
Magnitude of environment-recruitment relationships for 17 univariate analyses spanning nine species based on original (▪), extended (▴), and extended but balanced (•) time series. “Balanced” refers to estimates based on analyses where we randomly sub-sampled (1000 times) the pre-retest data so that there was an equal number of pre-and post-retest data points. Where there is more than one relationship per species-variable combination, letters indicate the source of data as being from the FMWT (A), Bay Study MWT (B), Bay Study OT (C), TNS (D for the overall Delta, and D* for Suisun Bay only), or Chinook salmon trawls (E1 at Chipps Island, E2 at Sacramento). The standardized effect is the slope of the relationship between abundance and the environmental variable under consideration in standard deviation units. For example, based on the updated time series, a one standard deviation unit increase in Flow at Freeport in February is expected to result in a 0.4 standard deviation unit increase in juvenile fall-run Chinook salmon index abundance in the Chipps Island trawl. Error bars are ± 1 SE.

In some instances, the addition of more years of data resulted in more variation in the relationship being explained (i.e., higher R^2^; Figure 5A, B). However, on average considering more recent data did not increase the estimated strength of the relationships, and in some cases reduced it (Figure 5A). For example, relationships between striped bass abundance and flow saw substantial declines in R^2^ with the addition of more years of data, despite the fact the magnitude of the relationship remained nearly identical (Figure 4). This occurred because the overall abundance of striped bass declined between the original time series and the updated one, which has been attributed to introduction of the Asian clam in 1987, but abundance still increased with increases in flow during both periods (Figure S1).

**Figure 5:**
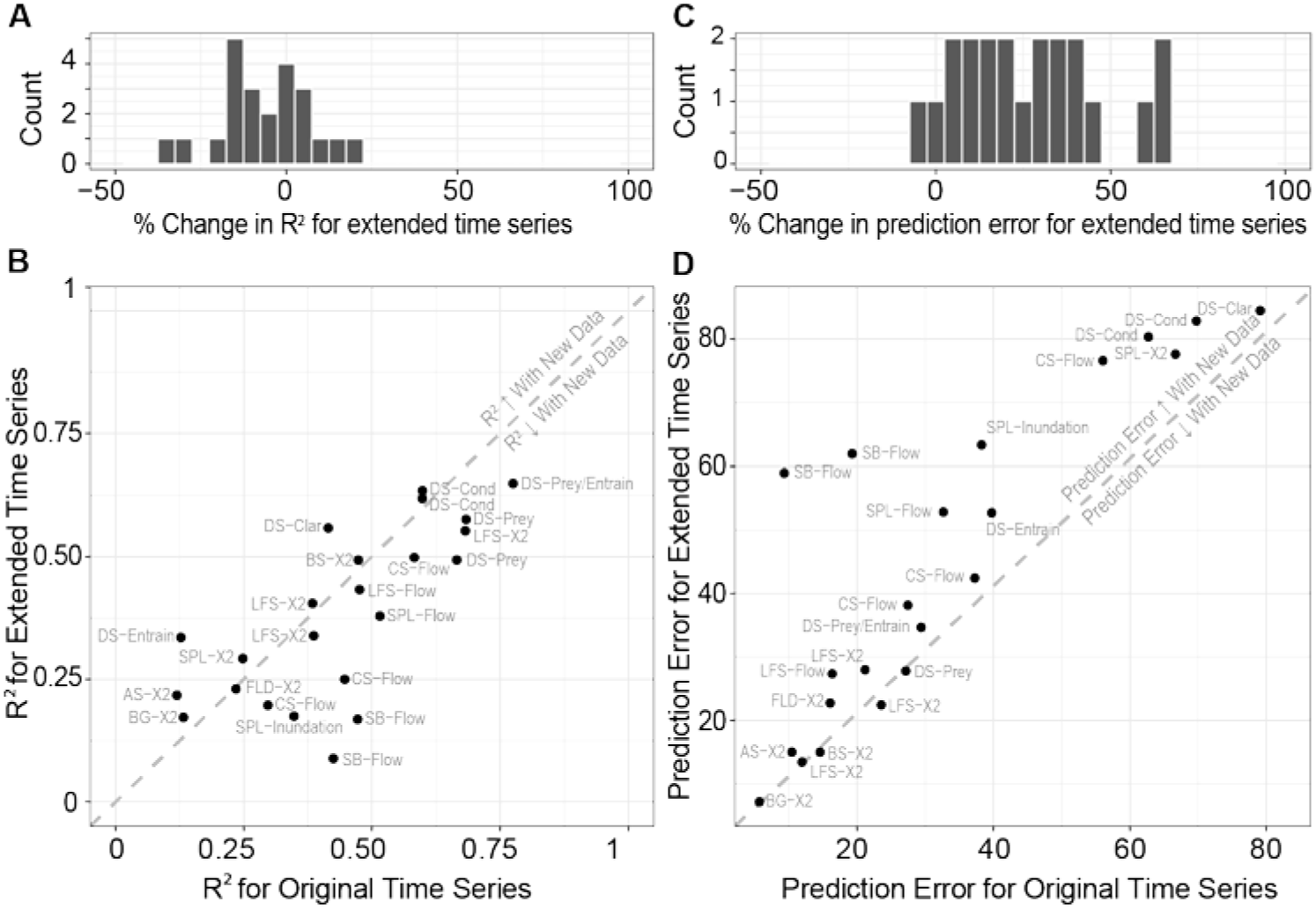
Percent change as well as absolute variation explained (R^2^; A, B) and predictive power (prediction error; C, D) of Bay Delta environment-recruitment relationships when additional years are added to the relationships. Estimates of percent change in prediction error for the striped bass – flow relationship are not included in C and D due to their very large values (~600%; see Table 1). Prediction error (see equation 1) is the percent error in future predictions relative to the average observed abundance for a given relationship. For example, a prediction error of 50% would mean that the relationship is expected to predict future abundances within +/- 25% of the mean predicted abundance. The first two letters of each label are species codes as follows: AS = American shad, BG = Bay goby, BS = Bay shrimp, DS = Delta smelt, LFS = longfin smelt, CS = Fall-run Chinook salmon, LFS = longfin smelt, SPL = Sacramento splittail, SF = starry flounder, SB = striped bass.

In most cases, considering more years of data did not improve the predictive power of the environment-recruitment relationships. Instead, counterintuitively, the prediction error of each relationship typically increased with the addition of more years of data (Figure 5C, D). The median percent increase in prediction error across the relationships retested was 30%. As with the variation explained, these changes were most pronounced for relationships between striped bass abundance and flow.

## Discussion

Our review of the literature identified 98 publications examining 420 individual environment-recruitment relationships in the Bay Delta. These relationships overwhelmingly focused on the influence of environmental variables on population abundance as opposed to other biological characteristics and were biased towards species that are either currently or historically listed as Threatened or Endangered. About half of these relationships are used in regulatory decision-making, but only one in five relationships have been retested to quantify the extent to which they stand the test of time. We retested 23 relationships using data that have accumulated in the years since they were first published. In contrast to Myers (1998), who found that the proportion of relationships that held up when retested was low, we found that when new data were used to retest previously published Bay Delta relationships, the direction and statistical significance of the relationships remained the same, though the amount of variation explained by the relationships and our ability to predict how a species will respond to change in their environment did not generally improve with more data. Instead, in most cases, prediction error actually increased when extending the time series, suggesting that accumulating more data will not necessarily improve the ability of these relationships to inform decision making (insofar as predictive power is useful for decision making). Perhaps this should not come as a surprise given the original relationships examined here were typically identified based on their ability to explain observed data (e.g., R^2^ from linear regression) as opposed their ability to predict future observations using, for example, approaches like data splitting and cross-validation (Power 1993; Harrel 2015). However, a large reduction in predictive power when relationships are re-tested with more data may be diagnostic of an established relationship that is breaking down (i.e., as is the case with striped bass and potentially Sacramento splittail in Figure 5), and might prompt action to search for unmeasured drivers of this change.

Our review of published environment-recruitment relationships in the Bay-Delta also highlights some methodological shortcomings of studies in this discipline. First, our review has made clear the great value of long-term data collection programs that follow standardized and consistent protocols to detect and validate long-term trends in biological variables, and has shown that a large share of studies in this space leverage these datasets. However, the availability and accessibility of such survey data may also reduce the likelihood that researchers in this region will embark on independent data collection to study other species and biological variables that are not the focus of existing surveys. Second, despite the accessibility of long-term survey data and the simplicity of analysis by correlation, we were still unable to reproduce originally published parameters for many relationships otherwise meeting our criteria for re-testing. In some cases, this was due to errors in the original work, unreported assumptions about data transformations that only became clear after contacting the authors, or the prior use of interpolated data that has since been corrected at the source by survey operators. These challenges highlight the importance of reproducibility in research in general, and into environment-recruitment relationships in particular, echoing a growing call for greater reproducibility both in ecology and across other scientific disciplines (Cassey and Blackburn 2006; Nosek et al. 2015; Borregaard and Hart 2016). Trends towards the use of coding for ecological analysis, and for the publication of that code alongside manuscripts as we do in our Supplementary Online Materials for this study, will play a significant role in improving the reproducibility of ecological research going forward (Mislan et al. 2016).

Our conclusions should be considered in light of potential biases in both our study selection criteria and in the type of data these studies draw upon. As with all studies based on literature review, our results are subject to publication bias (Cooper et al. 2009) relating to our decision to focus on peer-reviewed studies, a propensity towards publication of significant relationships in peer-reviewed journals, and a disproportionate number of publications on particular variables (e.g., X2 and flow) coming from a few very active authors in this field. In addition, recent work has shown that the long-term survey data used to create many of these relationships may itself be inherently biased by unquantified changes in detection probability. Detection probability, or catchability, may vary considerably over time with (1) overall abundance (i.e., it is more difficult to catch a rarer species or size class) (Mahardja et al. 2017), (2) with changing environmental conditions (e.g., catchability may decrease with increasing water clarity as fish are better able to see and avoid survey gear in clearer water) (Latour 2016), and (3) with differences in gear type across surveys (J. T. Peterson and M. Barajas, unpubl.). When not accounted for, these changes in detectability may be incorrectly interpreted as real changes in abundance or occupancy. Notwithstanding these potential biases, we believe that the breadth of species, environmental variables, and survey types covered in our analysis still allows us to draw some more general conclusions about the utility of environment-recruitment models, and to synthesize insights from the literature on best practices for the analysis, use, and refinement of environment-recruitment relationships to inform decision making in natural resource management.

### Correlation, causation, and strength of evidence

Correlations underpin most natural resource management decisions where one must predict how the environment responds to human action. These correlations are based on experience, and accumulated observations of the relationship between a species (e.g., abundance or survival) and its environment (e.g., flow, temperature, prey abundance, etc.). Such historic relationships are usually assumed to be causative, but we are often reminded of the adage “correlation does not equal causation”. Ideally, manipulative experiments can be used to determine whether a specific human action causes a response in an ecosystem component. Such learning by manipulation embraces the three key elements of experimental design: controls, randomization of treatments, and replication. Controls (i.e., monitoring systems that are not subject to a specific management intervention) provide critical contrast needed to disentangle the effects of the management actions (treatment) from other system change. Randomization ensures that the choice of which systems receive the management intervention is based on chance, thereby reducing the potential for confounding and bias due to factors not accounted for in the experimental design. Lastly, replication, whereby treatment and control groups are replicated over space and/or time, allows for more precise estimates of the effect of the management intervention.

Much has been learned in natural resource management through manipulative experiments. For example, hatcheries manipulate the timing and size of fish released to determine which combination result in optimal survival (e.g., Irvine et al. 2013) and hydroelectric facility operators manipulate timing and magnitude of flow releases to determine which flows are most likely to improve fish survival (e.g., Bradford et al. 2011). While manipulative experiments are the gold standard approach to establishing causation, scope for manipulative experiments decreases at increasing scales, and so we are left interpreting correlative relationships in order to manage some of the largest human-influenced ecosystems in the world.

When opportunities for learning by manipulation are limited or impossible, the weight of evidence for a hypothesized causal correlation can be assessed based on the strength, consistency, specificity and plausibility of the mechanism underlying the relationship (e.g., Hill 1965; Hilborn 2016). The stronger the association between two variables and implied magnitude of effect (e.g., a small [2%] versus large [50%] change in abundance), the more likely the relationship is causal. When the same relationship is found repeatedly across space (e.g., in other systems or species) and over time (e.g., with addition of more years of data), the stronger the evidence that it is indeed directly causal or related to the same causal driving variable (e.g., relationships to flows ultimately driven by precipitation) and not due to another unmeasured confounding factor. How likely are alternative explanations (hypotheses) for a given relationships? When few or no alternative explanations exist, causality is more likely than under circumstances where alternative explanations abound. Lastly, the plausibility of the mechanism underlying the association between two variables can shed light on the likelihood it is causal; when there is clear evidence of a mechanism that could be responsible for the relationship (e.g., from laboratory experiments) then there is stronger evidence that the relationship is indeed causal relative to instances where support for (or nature of) the underlying mechanisms is unknown.

While strength, consistency, specificity and plausibility can help guide the degree of support for a given relationship, they should not come at the cost of maintaining multiple working hypotheses, and evaluating the evidence for each simultaneously when using correlations to guide decision making (Hilborn and Mangel 1997; Plowright et al. 2008). An illustrative example of the simultaneous consideration of multiple working hypotheses is the development and application of a state-space multi-stage life cycle model to investigate for drivers of population decline in delta smelt by Maunder and Deriso (2011).

### Adaptive management in the face of ecosystem change

Even when relationships are truly causative, using past relationships to guide future management decisions can fail to have the intended effect when the system in which they occur changes over time (i.e., exhibits non-stationary). Such non-stationarity in aquatic systems can arise from both slow-moving environmental change or rapid regime shifts and “tipping points” between alternative stable states (Scheffer et al. 2001, 2009). There is widespread evidence of regime shifts in aquatic ecosystems arising from both natural (e.g., climate) and human (e.g., pollution, species introductions) caused factors (Carpenter 2003; Hunsicker et al. 2016) and the state of the ecosystem can have a strong influence on the outcomes of management actions. For example, large releases of hatchery salmon reduce the survival of endangered wild salmon but only during periods of poor ocean conditions (Levin et al. 2001), and translocation of wild juvenile salmon past hydropower installations carries greater benefits for their ocean survival in cooler but not warmer oceanic regimes (Gosselin et al. 2017). These examples highlight the fact that the benefits of management interventions (e.g., reducing hatchery releases to minimize impacts to wild fish, translocation of fish past barriers) are contingent upon the ocean regime the system is experiencing in any given year. In the Bay Delta, the introduction and rapid expansion of the invasive Asian clam in the late 1980s is believed to have caused a major increase in grazing pressure on phytoplankton, leading to a persistent decline in overall pelagic food resources (Carlton et al. 1990; Nichols et al. 1990; Baxter et al. 2010). The so-called “step change” towards this new stable state has had varying influences on different species within the ecosystem. For example, the regime shift due to Asian clam has led to a change in the overall magnitude (intercept) but not the rate of change (slope) of existing abundance-flow relationships for striped bass (Kimmerer et al. 2009, and reproduced in this study), but has been suggested to have driven a new abundance-flow relationship for threatened Delta smelt that brought with them significant implications for the way flows in this system are managed (CDFW 2016a,b).

Given the ubiquity of regime shifts and non-stationarities in aquatic systems (Möllmann and Diekmann 2012), including in the Bay Delta (Kimmerer 2002; Kimmerer et al. 2009; Thomson et al. 2010), how should one evaluate the evidence for environment-recruitment relationships and use them to inform decision making when system change is suspected? In some cases, change in a system will be so pronounced that there is little question about when it occurred and so the nature of a relationship post regime change should be the one that is used to inform future management actions. In other instances, the timing of abrupt change, and indeed whether it has occurred at all, will be uncertain for several years and so debated (at times vociferously) until clear evidence accumulates that a regime shift has occurred. In such cases, quantifying the statistical support for changes in system state using, for example, change-point analyses (Thomson et al. 2010) or dummy variables in linear regression (e.g., Kimmerer 2002; Kimmerer et al. 2009), is one way to quantitatively evaluate the evidence for system change. In addition, and perhaps more importantly, one should quantitatively evaluate the decision-making consequences of incorrectly assuming a regime shift has or has not occurred so as to be able to understand and clearly communicate the costs (e.g., biological, economic and social) of getting it wrong. Lastly, it has also been found that when non-stationarity is present (or suspected to be present), using recent observations to predict the consequences of alternative management actions can improve management outcomes (e.g., Ianelli et al. 2012).

In systems that have undergone dramatic change (e.g., tipping points), failure to regularly re-evaluate the durability and predictability of environment-relationships risks making management decisions based on information with increasingly large margins of error, with the potential for negative ecological, social and economic consequences. To illustrate this point we estimated prediction error for a few Bay delta environment-recruitment relationships spanning the period before and after the Asian clam invasion in 1987 for species that have (striped bass and longfin smelt; Kimmerer et al. 2009), and have not (Sacramento splittail; Kimmerer et al. 2009), responded to the invasion (Figure 6). For those species that declined in abundance coincident with the invasion, failure to account for this change results in relationships with increasing prediction error as time goes by after the invasion, relative to relationships that account for the change by including a step change in 1987 (striped bass and longfin smelt; Figure 6). In contrast, for Sacramento splittail, which did not appear to respond to the invasion, there is no benefit to including a step change in the relationship. Interestingly, these analyses suggest that the environment-recruitment relationships for striped bass and longfin smelt have experienced subsequent regime shifts (~1995 for striped bass and ~ 2005 for longfin smelt) that have further eroded their predictive power. This subsequent shift may be explained by observed changes in distribution likely to affect catchability in long-term surveys. Prior studies have suggested that a reduction on pelagic food resources due to overgrazing by Asian clam appears to have driven shifts in the distribution of young fish in the 1980s and 1990s away from the primary sampling regions of long-term surveys and towards areas characterized by fewer clams and better foraging prospects. This manifested as a lateral shift from deeper channel habitat preferentially sampled by annual surveys towards shallower slough habitat for striped bass (Sommer et al. 2011), and as a longitudinal shift from upstream habitat towards more saline downstream habitat for longfin smelt (Baxter et al. 2008; Sommer et al. 2011). Thus, these two shifts in prediction error align and illustrate with two stages rapid environmental change driven by a trophic cascade, the first likely related to a species introduction, and the second likely related to two different behavioural responses to adapt to the consequences of this introduction.

**Figure 6:**
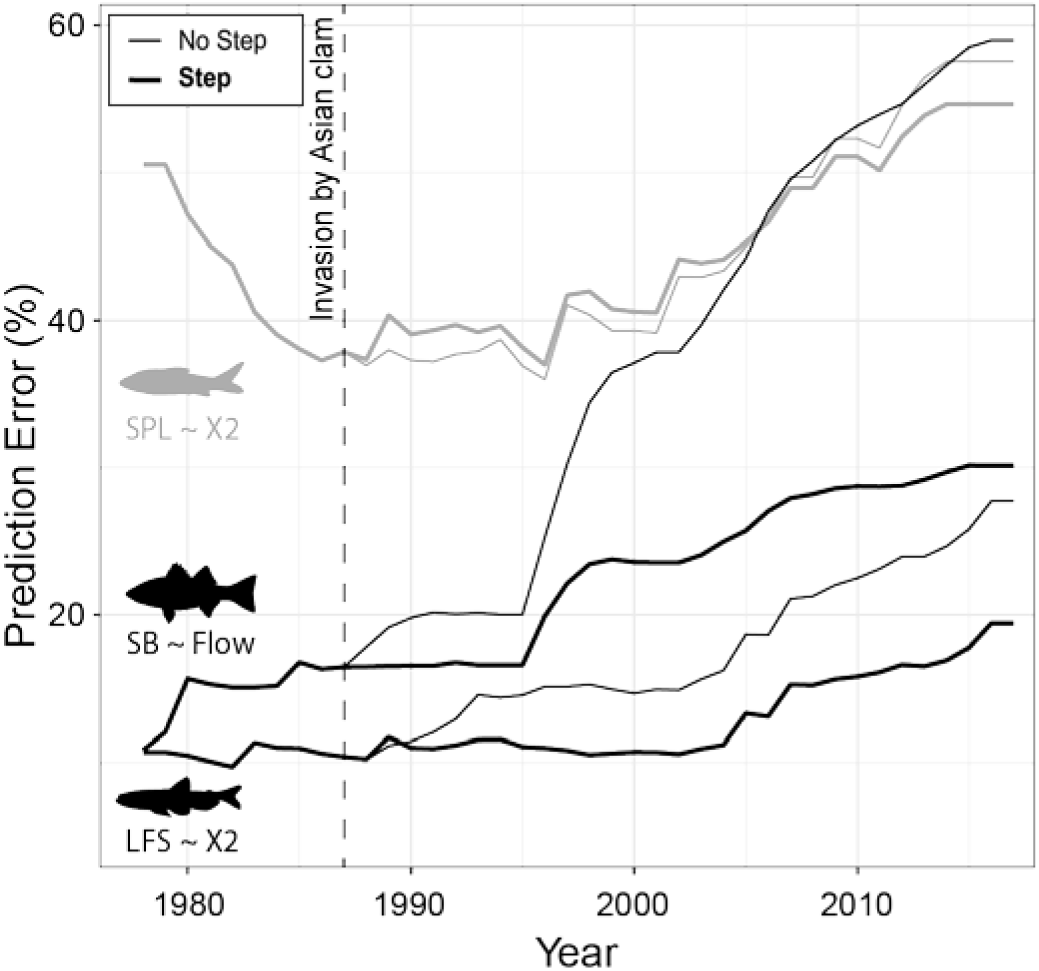
Prediction error (equation 1) relative to the logged index values used in the models over time for three illustrative Bay Delta environment-recruitment relationships modelled without (thin lines) and with (heavy lines) a step change to account for the regime shift associated with the introduction of the Asian clam circa 1987. Prediction error in each year is based on all years of data up to that point in time. Some species (black: SB - striped bass, LFS - longfin smelt) show strong responses to the regime shift and benefit from reduced prediction error with the inclusion of a step change in the models, while others (grey: SPL – Sacramento splittail) do not.

Our findings suggest that when environment-recruitment relationships underpin decision-making they should be re-evaluated on a regular basis as part of a broader adaptive management approach to ensure that they remain robust in the face of new data and continue to provide an accurate representation of a continually evolving ecosystem. Such an adaptive approach to evaluating environment-recruitment relationships is aligned with broader calls for increasing the implementation of more proactive adaptive management in Bay Delta ecosystems to address accelerating environmental change (Delta Independent Science Board 2015, 2016; Zandvoort et al. 2017).

### Environment-recruitment relationships in decision making

The widespread use, and at times misuse, of environment-recruitment relationships to inform decision making has produced several general insights into best practices for incorporating such relationships into natural resource management.

First, uncertainty should be both quantified and propagated in any analysis that seeks to predict the consequences of alternative management actions and identify those actions most likely to achieve desired objectives. This uncertainty comes in at least four distinct forms (e.g., Peterman 2004): (1) natural variation in both physical and biological processes, (2) uncertainty due to imperfect assessment arising from measurement error, (3) structural uncertainty due to incomplete understanding of how a system functions, and lastly (4) outcome uncertainty or implementation error in how well a given management target (e.g., increase flow by 20%) is achieved by a management action (e.g., releasing water from a reservoir). While uncertainty resulting from points 2-4 can in theory be reduced by improved measurements, greater understanding of system function, and better management control, all of which can be accomplished to some extent with the collection of more years of data, uncertainty arising from natural variability is irreducible. We found that prediction error was not reduced for Bay Delta environment-recruitment relationships with the accumulation of more years of data, and in fact increased in many cases. This finding suggests that natural variation in the physical or biological processes of the system is an important and ongoing source of uncertainty that will limit the extent to which improvements in predictions of how taxa from the Bay Delta will respond to changing environmental conditions and human action can be achieved.

Second, there is increasing recognition of the value of developing and using life cycle models to evaluate the predicted consequences of alternative management actions on species of concern in the face of this uncertainty (e.g., Good et al. 2007; Ruckelshaus et al. 2002; Zeug et al. 2012). In contrast to single life stage, habitat type, or environmental relationships, life-cycle models simultaneously consider extrinsic (environment, management action) and intrinsic (density dependence) factors influencing multiple life stages. Life cycle models can either be mechanistic where survival between life stages is based on specific mechanisms (Scheurell et al. 2006) or statistical where life stage specific survival is not defined by specific mechanistic relationships (Nobriga and Rosenfield 2016). The use of life cycle models allows for more realistic and comprehensive evaluation of the predicted outcomes of alternative management actions than considering single life stage, habitat type, or environmental relationships, because it considers environmental effects across linked stages in a life cycle while also accounting for population processes (e.g., growth, movement, mortality and reproduction).

Even when uncertainty is successfully incorporated into modelling approaches, the broader question remains – how can we account for and propagate uncertainty into the broader management of a complex system with many conflicting management objectives when our understanding of that system is not, and will never be, complete? In a review of management approaches in the Bay-Delta ecosystem for the National Research Council, the Committee on Sustainable Water and Environmental Management proposes that agencies should adopt management approaches that assume “ ‘universal nonstationarity,’ or the idea that all aspects of the environment will constantly be changing.” (NRC 2012). Such approaches may prove challenging for many traditional decision-making pathways, which are often constrained by static or slow-moving policy frameworks (Aladjem 2013; Delta Independent Science Board. 2016). However, a number of approaches with shared characteristics have been developed to help facilitate formalized decision-making in complex systems characterized by their uncertainty and are among the approaches recommended by the NRC review. Among these are decision analysis or decision scaling, risk assessment, and management strategy evaluation.

Decision analysis, a form of risk assessment, is a systematic approach to incorporating uncertainties in the managed system into decision making using models to project outcomes of alternative actions (Peterman and Anderson 1999). These outcomes (or performance measures) act as criteria for comparing and ranking the actions against specific objectives (e.g., ecological, social and economic). Decision analysis can be coupled with stochastic simulations (e.g., Peters and Marmorek 2001) to identify management “actions” that explicitly consider multiple sources of uncertainty, where management actions refer to a combination of sampling design for data collection, approaches to analyzing the data, and decision rules for how to manage a specific aspect of the system. However, such risk assessments and decision analysis can overestimate risk by failing to account for management reactions to the information provided by future data. An alternative version of the approach known as “decision scaling” begins with a bottom-up analytic framework that first identified the most critical management uncertainties that influence decision-making, and uses this information to tailor top-down environmental monitoring and modelling to provide the most credible information at appropriate temporal and spatial scales to project the outcomes of alternative actions (e.g., different flow management regimes) under a range of possible futures (e.g., wet and dry years) (NRC 2012). Explicitly considering management feedback is also a central feature of what has been variously referred to as closed-loop simulations (Walters 1986), Management Strategy Evaluations (Sainsbury et al. 2000), and Management Procedure Evaluations (Butterworth and Punt 1999), which are in effect risk assessment methods and formal decision analyses which consider management feedback. While most widely applied in relation to fisheries and cetacean conservation and management, Management Strategy Evaluations have also been used to inform decision making focused on multi-species and ecosystem objectives (reviewed by Punt et al. 2014). All of these approaches lend themselves to robust evaluation of the value of different sources of information, including environment-recruitment relationships and the uncertainty inherent therein, into a broader assessment of the best way forward for natural resource management decisions in complex systems.

### Conclusions

Moving forward, there is a growing recognition of the importance of maintaining multiple working hypotheses when quantifying the support for correlations in environmental management (Hilborn 2016), that quantitative assessment of policies that consider these relationships should be done using approaches that allow for realistic incorporation and propagation of multiple sources of uncertainty (Peterman 2004), and that, ultimately, managers in the Bay-Delta and elsewhere should identify policies that are robust to a range of alternative hypotheses (NRC 2012; Schindler and Hilborn 2015).

Despite advances in the tools available to improve our assessment of environment-recruitment relationship and their consideration in decision making, we should remain humble in our zeal to either accept them as fact or discount them entirely because they are “just correlations”. As Hill emphasized in his 1965 Presidential Address on correlation and causation to the Royal Society of Medicine (Hill 1965): “All scientific work is incomplete – whether it be observational or experimental… [and] is liable to be upset or modified by advancing knowledge. That does not confer upon us a freedom to ignore the knowledge we already have, or to postpone the action that it appears to demand at a particular time.”

## Acknowledgements

This work was supported through funding from the Metropolitan Water District of Southern California to authors NT and BMC. We would like to thank Ted Sommer, Shawn Acuña, Will Satterthwaite, Alison Collins, and Rebecca Sheehan for their help in identifying additional environment-recruitment relationships; B.J. Miller, Matt Nobriga, Fred Feyrer, Shawn Acuña, and Ted Sommer for their assistance in re-creating the original relationships from their published studies; B.J. Miller, Kathy Hieb of the California Department of Fish and Wildlife San Francisco Bay Study, and Derick Louie of the California Department of Water Resources for their assistance in obtaining raw data needed for re-testing relationships; and F. Poulson for his assistance in deriving some of the required abundance indices from raw data.

